# Mutations in *C11ORF70* cause primary ciliary dyskinesia with randomization of left/right body asymmetry due to outer and inner dynein arm defects

**DOI:** 10.1101/218206

**Authors:** Inga M. Höben, Rim Hjeij, Heike Olbrich, Gerard W. Dougherty, Tabea Menchen, Isabella Aprea, Diana Frank, Petra Pennekamp, Bernd Dworniczak, Julia Wallmeier, Johanna Raidt, Kim Gjerum Nielsen, Maria C. Philipsen, Francesca Santamaria, Laura Venditto, Israel Amirav, Freerk Prenzel, Kaman Wu, Miriam Schmidts, Niki T. Loges, Heymut Omran

## Abstract

Primary ciliary dyskinesia (PCD) is characterized by chronic airway disease, male infertility and randomization of the left/right body axis caused by defects of motile cilia and sperm flagella. We identified loss-of-function mutations in the open reading frame *C11ORF70* in PCD individuals from five distinct families. Transmission electron microscopy analyses and high resolution immunofluorescence microscopy demonstrate that loss-of-function mutations in *C11ORF70* cause immotility of respiratory cilia and sperm flagella, respectively, due to loss of axonemal outer (ODAs) and inner dynein arms (IDAs), indicating that C11ORF70 is involved in cytoplasmic assembly of dynein arms. Expression analyses of *C11ORF70* showed that *C11ORF70* is expressed in ciliated respiratory cells and that the expression of *C11ORF70* is upregulated during ciliogenesis, similar to other previously described cytoplasmic dynein arm assembly factors. Furthermore, C11ORF70 shows an interaction with cytoplasmic ODA/IDA assembly factor DNAAF2, supporting our hypothesis that C11ORF70 is a novel preassembly factor involved in the pathogenesis of PCD. The identification of a novel genetic defect that causes PCD and male infertility is of great clinical importance as well as for genetic counselling.

---

Cilia are hair-like organelles extending from nearly all types of polarized cells. Motile cilia in distinct cell types in the human body perform essential biological functions such as generation of fluid flow and mucociliary clearance of the airways^(1)^. The basic structure of motile cilia consists of a ring of nine peripheral microtubule doublets, which surround one central pair (9+2 structure). The peripheral ring is connected to the central pair (CP) through radial spokes (RS) and the nexin-dynein regulatory complex (N-DRC). The CP, the N-DRC and the inner dynein arms (IDAs) are responsible for modulation and regulation of the ciliary beating^(2,3)^ while the outer dynein arms (ODAs) are responsible for the beat generation. ODAs and IDAs are large multimeric protein complexes that are pre-assembled in the cytoplasm before transport to the axonemes^(4)^. There are at least two types of ODAs in humans: type 1 containing the axonemal dynein heavy chains (HC) DNAH5 and DNAH11, located proximally, and type 2 containing the dynein HCs DNAH5 and DNAH9, located distally in the ciliary axonemes^(5,6)^. In *Chlamydomonas reinhardtii*, there are seven distinct IDA complexes that can be divided in three groups: IDA group I1, I2 and I3^(7,8)^. The IDA complex of group I1 contains two HCs, while the three IDA complexes of group I2 contain each one HC that associates with the dynein light chain p28. The IDA complexes of group I3 also contain each one HC which associates with centrin. The identification of proteins responsible for correct assembly and the composition of these protein complexes are critical to understanding the disease mechanisms of motile cilia-related disorders such as primary ciliary dyskinesia (PCD).

Primary ciliary dykinesia (PCD, MIM 244400) is a rare genetic disorder caused by immotile or dyskinetic cilia, with prevalence in the range of 1:4.000 to 1:20.000^(9)^. Ciliary dysfunction in upper and lower airways leads to defective mucociliary clearance of the airways and subsequently to recurrent airway inflammation and bronchiectasis (Figure 1) and progressive lung failure. Dysfunction of cilia of the left-right organizer (LRO) present during early embryonic development results in randomization of the left/right body asymmetry. Approximately half of the PCD patients exhibit *situs inversus totalis* (Figure 1), referred to as Kartagener’s syndrome^(9)^. More rarely, other situs anomalies associated with complex congenital heart disease are observed^(10)^. Defects underlying motile cilia dysfunction often affect sperm flagella function and cause male infertility in PCD individuals, warranting assisted reproductive technologies. Another consequence of ciliary dysfunction, particularly evident in PCD mouse models, is hydrocephalus formation caused by disrupted flow of the cerebrospinal fluid through the cerebral aqueduct connecting the third and fourth brain ventricles^(11)^. Although dysmotility of the ependymal cilia is not sufficient for hydrocephalus formation in humans, probably due to morphological differences between the mouse and human brain, the incidence of hydrocephalus, secondary to aqueduct closure, is increased in PCD individuals^(11)^. PCD diagnosis is difficult and relies on a combination of tests including nasal nitric oxide (nNO) production rate measurement, high-speed video microscopy analysis of cilia and ciliary structure analyses by transmission electron microscopy (TEM)^(12)^ and high-resolution immunofluorescence microscopy.

**Figure 1.**
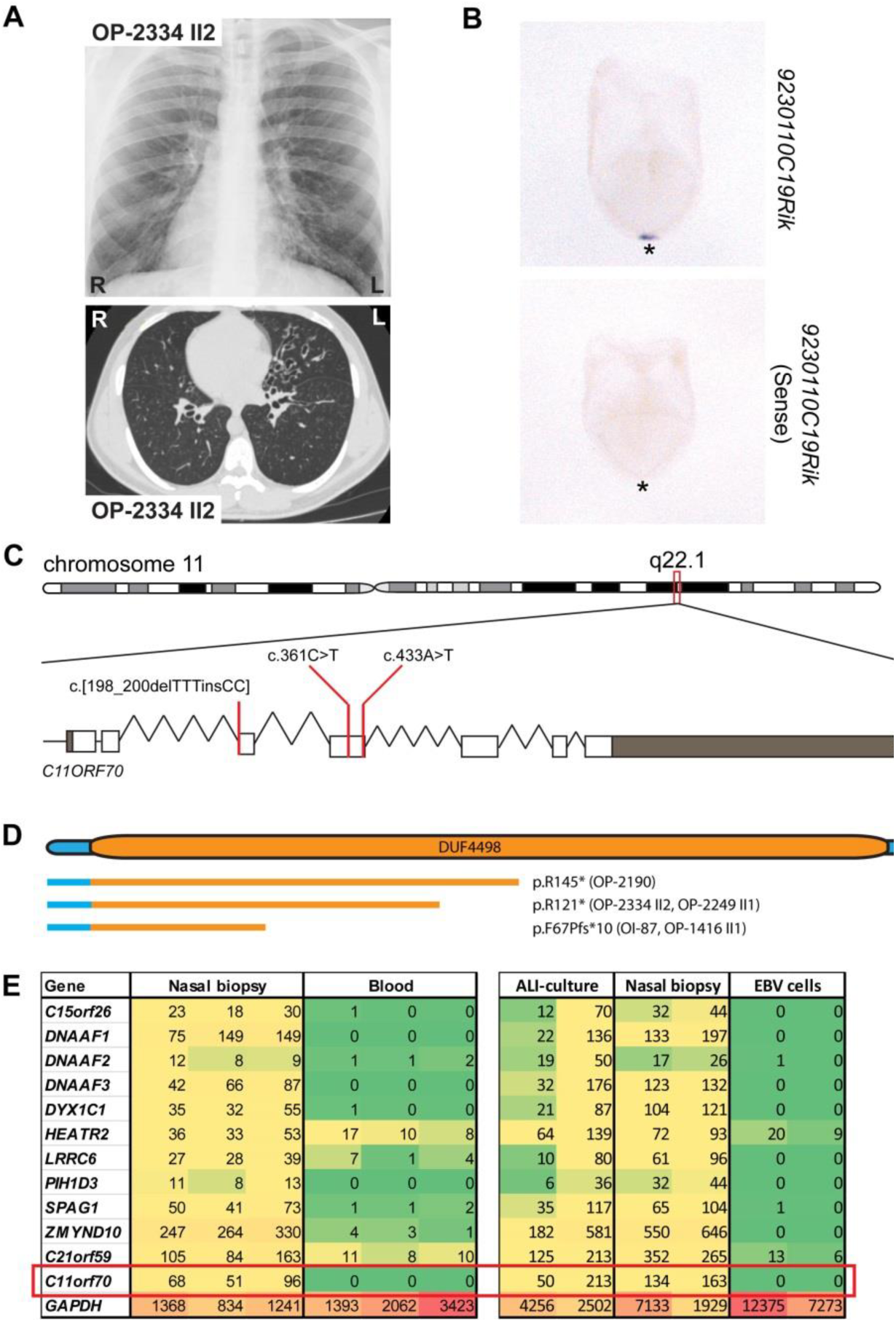
Recessive loss-of-function mutations in *C11ORF70* cause primary ciliary dyskinesia with randomization of left/right body asymmetry. (A) The chest X-ray radiograph of PCD individual OP-2334 II2 depicts *situs inversus totalis*. The computed tomography scan of OP-2334 II2 shows chronic airway disease with bronchiectasis in the middle lobe and mucus plugging. (B) *In situ* hybridization analyses of wildtype mouse 8.25 dpc (days post coitum) mouse embryos reveal expression of *9230110C19Rik* (*C11orf70 ortholog*) exclusively at the left/right organizer (frontal view; asterisks in the ventral position of the left/right organizer). By contrast, the negative control utilizing the sense probe does not show any staining. (C) Schematic presentation of chromosome 11 and the exon-intron structure of *C11ORF70* with untranslated (gray) and translated (white) regions. The positions of the three identified mutations in the five unrelated families are indicated by red lines. (D) Protein model of C11ORF70 with the domain of unknown function 4498 (DUF4498) and the predicted truncated proteins as consequences of the three *C11ORF70* loss-of-function mutations. (E) Differentiation-specific and tissue-specific expression profiles of known genes encoding proteins involved in outer and inner dynein arm assembly and *GAPDH* as a housekeeping gene. Raw RNA-seq data were normalized against *PPIH* (*peptidylprolyl isomerase H*). The preassembly factor genes have the strongest expression in native material of nasal brushing biopsies. Comparable expression profiles are observed in ALI-cultured epithelial nasal cells and in nasal brushing biopsies. EBV-cells (immortalized lymphocytes) and blood cells show no or weak expression of genes encoding cytoplasmic dynein assembly factors. *GAPDH* shows a high expression in all analyzed tissues.

Until now, mutations in 37 genes have been linked to PCD, responsible for an estimated 70% of cases^(13)^. Genetic analyses of PCD individuals identified several autosomal recessive mutations in genes encoding for axonemal subunits of the ODA and ODA-docking complexes^(14–26)^, the N-DRC^(27–30)^ and the 96-nm axonemal ruler proteins^(31,32)^, the RS^(33–36)^ and CP associated proteins^(37,38)^ (Figure 2). Although the process of cytoplasmic pre-assembly of dynein arms is still poorly understood, several genes encoding proteins involved in this process were identified: *DNAAF1* (*LRRC50*, MIM 613190)^(39,40)^, *DNAAF2* (*KTU*, MIM 612517)^(41)^*, DNAAF3* (*C19orf51*, MIM 614566)^(42)^, *DNAAF4* (*DYX1C1*, MIM 608709)^(43)^, *DNAAF5* (*HEATR2*, MIM 614864)^(44)^, *LRRC6* (MIM 614930)^(45)^, *ZMYND10* (MIM 607070)^(46,47)^, *SPAG1* (MIM 603395)^(48)^ and *C21ORF59* (MIM 615494)^(49)^. Three X-linked PCD variants have been reported so far, one caused by mutations in *PIH1D3* (MIM 300933)^(50,51)^ resulting in absence of ODA and IDA and two PCD variants associated with syndromic cognitive dysfunction or retinal degeneration caused by mutations in *OFD1* (MIM 311200) and *RPGR* (MIM 312610), respectively^(52,53)^. Here, we describe a new ODA/IDA defect caused by recessive loss-of-function mutations in the open reading frame of *C11ORF70*.

**Figure 2.**
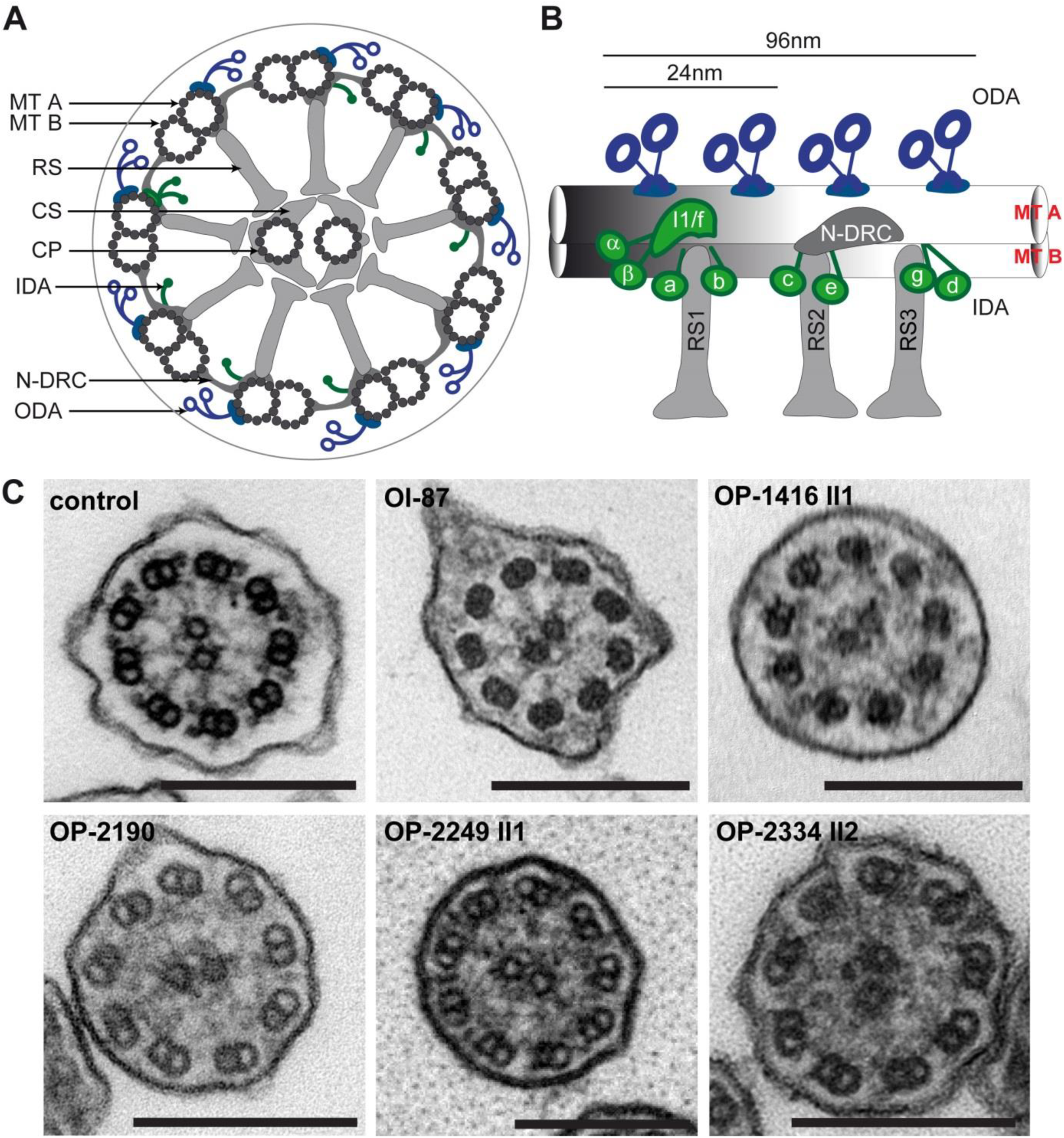
*C11ORF70*-mutant respiratory ciliary axonemes exhibit ultrastructural defects of the outer and inner dynein arms. (A) Schematic diagram of a cross-section through a motile respiratory 9+2 cilium. (B) Arrangement of ODAs and IDAs within the 96-nm unit in human ciliary axonemes. (C) Transmission electron microscopy photographs of cross-sections through respiratory epithelial cilia demonstrate absence of ODAs and IDAs in PCD individuals OI-87, OP-1416 II1, OP-2190, OP-2249 II1 and OP-2334 II2 carrying biallelic *C11ORF70* mutations when compared to a control cilium. Scale bar represents 200 nm. CP, central pair; CS, central sheath; IDA, inner dynein arm; MT A, microtubule A; MT B, microtubule B; N-DRC, nexin-dynein regulatory complex; ODA, outer dynein arm; RS, radial spoke.

We performed targeted-exome sequencing in 15 PCD individuals with combined ODA and IDA defects of unknown genetic cause. Signed and informed consent was obtained from individuals fulfilling diagnostic criteria of PCD^(54)^ and from family members using protocols approved by the Institutional Ethics Review Board at the University of Muenster. Genomic DNA was isolated by standard methods directly from blood samples. Targeted-exome sequencing of genomic DNA was performed at the Cologne Center for Genomics (CCG). For enrichment, the NimbleGen SeqCap EZ Human Exome Library v2.0 was used. Enriched preparations were sequenced with the HiSeq2000 platform (Illumina) as paired end 2 x 100 base pairs reads. The 30x coverage was in the range of 79-86%. The genome sequence hg38 was used as reference to map sequencing reads that passed quality filtering. Variants that were present in the dbSNP database, the 1000 Genomes Project polymorphism and the Genome Aggregation Database (gnomAD) with a minor allele frequency >0.01 were excluded. We focused on nonsynonymous mutations, splice-site substitutions and indels following an autosomal-recessive inheritance pattern. We identified in five PCD individuals loss-of-function mutations in *C11ORF70*. In OP-1416 II1 and OI-87, we identified a homozygous deletion of three thymine residues and an insertion of two cytosine residues (c.[198_200delTTTinsCC]) resulting in a frame shift and predicted premature stop of translation (p.F67Pfs*10; Figure 1 and Figure S1). In OP-2249 II1 and OP-2334 II2 we identified a transition from C>T at position 361, and in OP-2190 II1 a transversion from A>T at position 433 both resulting in stop codons (p.R121* and p.R145*; Figure 1 and Supplemental Figure S1).

All individuals with loss-of-function mutations in *C11ORF70* show classical PCD symptoms (Table 1) such as chronic sinusitis, chronic otitis media, and chronic lower respiratory tract infections as well as bronchiectasis in the middle lobe and mucus plugging (shown for OP-2334 II2 in Figure 1). Four of the five affected individuals had neonatal respiratory distress syndrome. In addition, one individual has *situs inversus totalis*, consistent with randomization of left/right body asymmetry (Figure 1). The nasal nitric oxide production rate measured with the Niox Mino (Aerocrine) or EcoMedics CLD88 (EcoMedics) was low in all affected individuals (Table 1). High-speed videomicroscopy analyses of ciliary motility in nasal respiratory epithelial cells showed completely immotile cilia (Supplemental Video 1, 2 and 3) in contrast to control cilia (Supplemental Video 4). Ciliary beat frequency and beating pattern was assessed with the SAVA system^(55)^. Respiratory epithelial cells were analysed with a Zeiss AxioVert A1 microscope (40x and 63x phase contrast objective) equipped with a Basler sc640-120fm monochrome high-speed video camera (Basler, Ahrensburg, Germany) set at 120 frames per second. The ciliary beating pattern was evaluated on slow-motion playbacks. Two individuals (OI-87 and OP-2334 II2) reported fertility problems. OI-87 gave birth to one child after in vitro fertilization. OP-2334 II2 underwent fertility testing in the past and was diagnosed with moderate oligozoospermia (reduced number of sperms) and severe asthenozoospermia (immotility of sperm flagella) (Table S1). TEM analyses of respiratory cilia isolated from individuals with biallelic *C11ORF70* mutations and sperm flagella from OP-2334 II2 displayed both ODA and IDA defects in all cases (Figure 2 and Figure 5). We performed high-speed video microscopy analyses of sperm flagella from OP-2334 II2 and in contrast to control (Supplemental Video 5), sperm cells from OP-2334 II2 were completely immotile (Supplemental Video 6 and 7) consistent with a loss of dynein arms. TEM of human respiratory cilia and sperm flagella was performed as previously described^(37)^.

**Table 1.** Clinical findings of PCD individuals with mutations in *C11ORF70*.

| Subject | Sex | SI | nNo<br>[nl/min] | Neonatal<br>RDS | Recurrent<br>Pneumonia | Recurrent<br>Respiratory<br>Infections | Bronchiectasis | Chronic<br>Wet<br>Cough | Otitis<br>Media | Hearing<br>Disorder | HVMA | Congenital<br>Heart<br>Defect | Fertility<br>Defect |
| --- | --- | --- | --- | --- | --- | --- | --- | --- | --- | --- | --- | --- | --- |
| OI-87 | F | no | 26 | yes | yes | yes | yes | yes | yes | yes | n.a. | no | probably<br>(one child<br>after IVF) |
| OP-1416 II1 | F | no | 13.3 | yes | yes | yes | yes | n.a. | n.a. | yes | immotile | no | probably |
| OP-2190 | F | no | 5 | yes | n.a. | yes | yes | yes | yes | n.a. | n.a. | no | n.a. |
| OP-2249 II1 | F | no | 5.7 | no | n.a. | yes | yes | yes | yes | n.a. | immotile | no | n.a. |
| OP-2334 II2 | M | yes | 28 | n.a. | n.a. | yes | yes | yes | yes | yes | immotile | no | yes |

Abbreviations: F, Female; M, Male; SI, Situs inversus; nNO, nasal Nitric Oxide; RDS, Respiratory Distress Syndrome; HVMA, High-speed video microscopy analyses; n.a., not available; IVF, in vitro fertilization

Next we analyzed *C11ORF70* expression in various tissues to test whether the expression is consistent with function in motile cilia and the observed clinical phenotype. Therefore, we studied tissue-specific expression of *C11ORF70* in EBV-transduced lymphocytes, nasal brushing biopsies and respiratory cells grown on air liquid interface (ALI)-culture to full differentiation, each obtained from two healthy controls. Human transcriptome profiles were generated by next generation sequencing. RNA from nasal epithelial cells, ALI cultured epithelial cells and EBV-infected lymphocytes were isolated using the RNeasy Mini Kit. Next-Generation-Sequencing IonSphere positive particles were generated using the IonOneTouch™ System 2 (Thermo Fisher). Prepared IonSphere positive particles were loaded into an Ion 318™ Chip (Thermo Fisher) and sequenced using the Ion Proton™ platform for Next-Generation-Sequencing. Sequenced amplicons were aligned against the reference transcriptome hg19_AMpliSeq_Transcriptome_ERCC_v1.fastq (for detailed description of ALI cell culture and human transcriptome generation see also Supplemental Material and Methods). Output data of a small selection of genes were normalized against *peptidylprolyl isomerase H* (*PPIH*). Mutations in genes encoding dynein axonemal assembly factors (DNAAFs) result in combined ODA and IDA defects resemble findings observed in *C11ORF70* mutant cilia. Similar to known genes encoding cytoplasmic dynein preassembly factors, *C11ORF70* was strongest expressed in native material of nasal brushing biopsies. Comparable expression was observed in nasal brushing biopsies and ALI-cultured nasal epithelial cells, whereas EBV-infected lymphocytes and blood cells usually show no or weak expression levels (Figure 1). Interestingly, expression of *C11ORF70* was upregulated during ciliogenesis between day 3 and day 15, similar to the expression of genes encoding DNAAFs involved in ODA/IDA assembly (Figure S2). To determine if loss of *C11ORF70* function could explain the *situs* abnormalities observed in one of the investigated PCD individuals with biallelic mutations, we performed *in situ* hybridization analyses of gastrulating mouse embryos (for further details see also Supplemental Material and Methods) during the developmental period when the LRO is present and body laterality is determined. We detected expression of *9230110C19Rik*/*C11orf70* during this essential developmental stage. Interestingly, the expression was restricted exclusively to the left/right organizer (Figure 1 and Figure S3). This finding strongly indicates that C11ORF70 is involved in function of motile nodal monocilia and that deficiency of this protein probably causes an impairment of nodal ciliary beating, resulting in randomization of the left/right body axis.

To further understand the defect at the molecular level, we performed high resolution immunofluorescence (IF) microscopy of respiratory cilia using antibodies targeting components of the ODAs (DNAH5, DNAH9, DNAH11, DNAI1 and DNAI2), IDAs (DNALI1, IDA group I2 and DNAH6, IDA group I3) and ODA docking complexes (TTC25). High resolution IF was performed as previously described (42). We previously showed that respiratory cilia contain two distinct ODA types: type 1, which contains the dynein heavy chains DNAH5 and DNAH11 (proximal ciliary axoneme), and type 2, which contains the dynein heavy chains DNAH5 and DNAH9 (distal ciliary axoneme) (Figure 3)^(5,6)^. The ODA components DNAH5, DNAH9, DNAI1 and DNAI2 were absent or severely reduced from the respiratory ciliary axonemes of the affected individuals (Figure 3 and Figures S4, S5, S6 and S7). Interestingly, DNAH11 localization at the proximal part of the axoneme was not altered in *C11ORF70* mutant cilia (Figure S8). However, TTC25 localization along the ciliary axoneme was not altered (Figure S9), indicating that loss of function of *C11ORF70* does not affect composition and assembly of the ODA docking complex. We next analyzed if assembly of the IDAs is also disturbed by C11ORF70 deficiency. We found that DNALI1 and DNAH6 were severely reduced or completely absent from the ciliary axonemes in these PCD individuals (Figure 4 and Figures S10 and S11), indicating that C11ORF70 is also involved in the assembly of the IDA-group I2 and I3. Because OP-2334 II2 was diagnosed with severe asthenozoospermia and TEM analyses demonstrated a combined ODA/IDA defect, we performed high resolution IF of sperm flagella using antibodies directed against DNAI1, DNAI2 and DNALI1. Both ODA components DNAI1 and DNAI2 as well as the IDA component DNALI1 were absent or severely reduced from flagellar axonemes (Figure 5), demonstrating that C11ORF70 is also involved in the preassembly of ODAs and IDAs in sperm cells.

**Figure 3.**
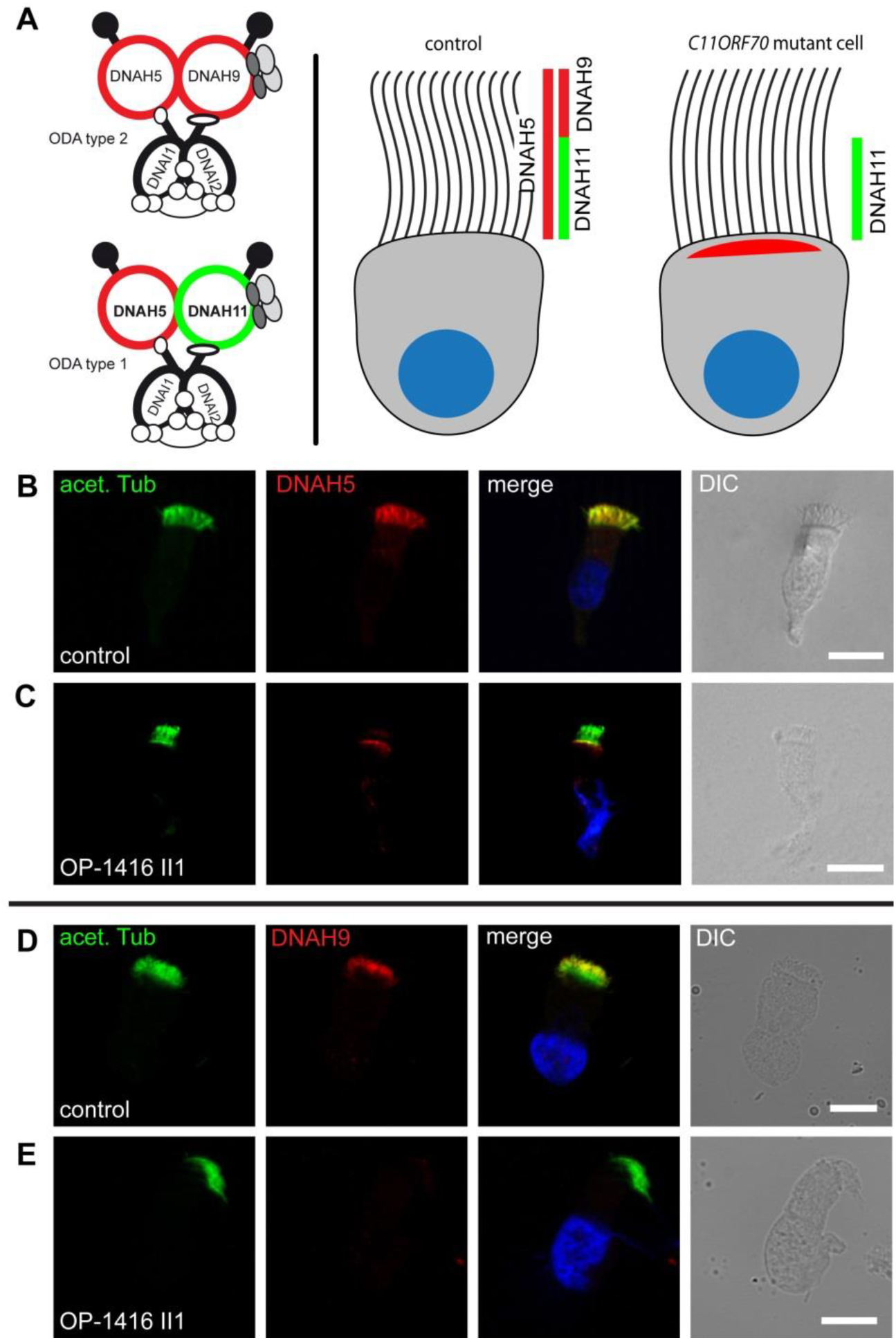
Absence of the outer dynein arm heavy chains DNAH5 and DNAH9 in respiratory cilia of PCD patients carrying *C11ORF70* mutations. (A) Schematics of ODAs type 1 and type 2 show that DNAH11 is a component of ODA type 1 and localizes only to the proximal part, while DNAH9 is a component of the ODA type 2 and localizes only to the distal part of the ciliary axonemes. DNAH5 is a component of both ODA types and localizes throughout the ciliary axoneme. In *C11ORF70*-mutant cilia, DNAH5 and DNAH9 are absent from the ciliary axonemes, whereas DNAH11 localization is not affected by mutations in *C11ORF70* (Figure S8). (B and C) Respiratory cilia double-labeled with antibodies directed against acetylated α-tubulin (green) and DNAH5 (red) show colocalization of DNAH5 with acetylated α-tubulin along the cilia from unaffected controls (B, yellow). In contrast, DNAH5 is absent or severely reduced in *C11ORF70*-mutant axonemes (C, shown for OP-1416 II1). (D and E) Respiratory epithelial cells from control and PCD individuals were double-labeled with antibodies directed against acetylated tubulin (green) and DNAH9 (red). Acetylated α-tubulin localizes to the entire length of the ciliary axoneme, whereas DNAH9 localization is restricted to the distal part of the axoneme in healthy control cells (D). In *C11ORF70*-mutant cells, DNAH9 is absent from the ciliary axonemes (E, shown for OP-1416 II1). Nuclei were stained with Hoechst33342 (blue). Scale bars represent 10 μm.

**Figure 4.**
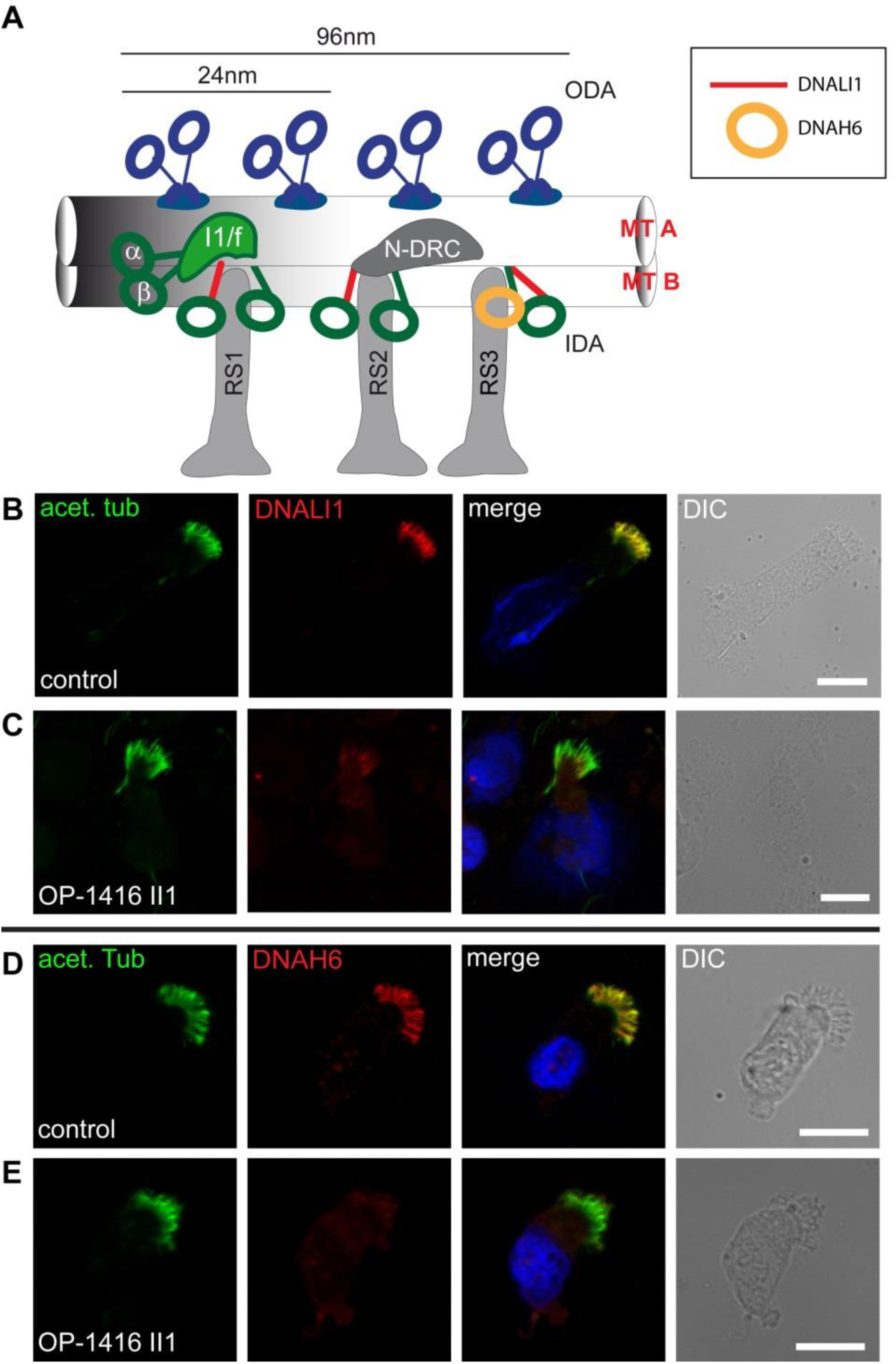
In *C11ORF70*-mutant respiratory cilia the inner dynein arm light chain DNALI1 (component of the IDA-group I2) as well as the inner dynein arm heavy chain DNAH6 (component of the IDA-group I3) are absent. (A) Schematic of the arrangement of ODAs (blue) and IDAs (green) within the 96-nm unit in the human ciliary axoneme. IDA complexes a, c and d belong to the IDA-group I2 characterized by the presence of the dynein light chain DNALI1 (red lines). IDA complex g, contains the dynein heavy chain DNAH6 (yellow circle), and belongs to the IDA-group I3. (B and C) Respiratory cilia were double-labeled with antibodies directed against acetylated α-tubulin (green) and DNALI1 (red) show colocalization of DNALI1 with acetylated α-tubulin along the cilia from unaffected controls (B, yellow). In contrast, DNALI1 is absent or severely reduced in *C11ORF70-*mutant axonemes (C, OP-1416 II1). (D and E) Respiratory epithelial cells from control and PCD individuals were double-labeled with antibodies directed against acetylated α-tubulin (green) and DNAH6 (red). DNAH6 colocalizes with acetylated tubulin along the cilia from unaffected controls (D, yellow). In contrast, DNAH6 is absent or severely reduced in *C11ORF70*-mutant axonemes (E, OP-1416 II1). Nuclei were stained with Hoechst33342 (blue). Scale bars represent 10 μm.

**Figure 5.**
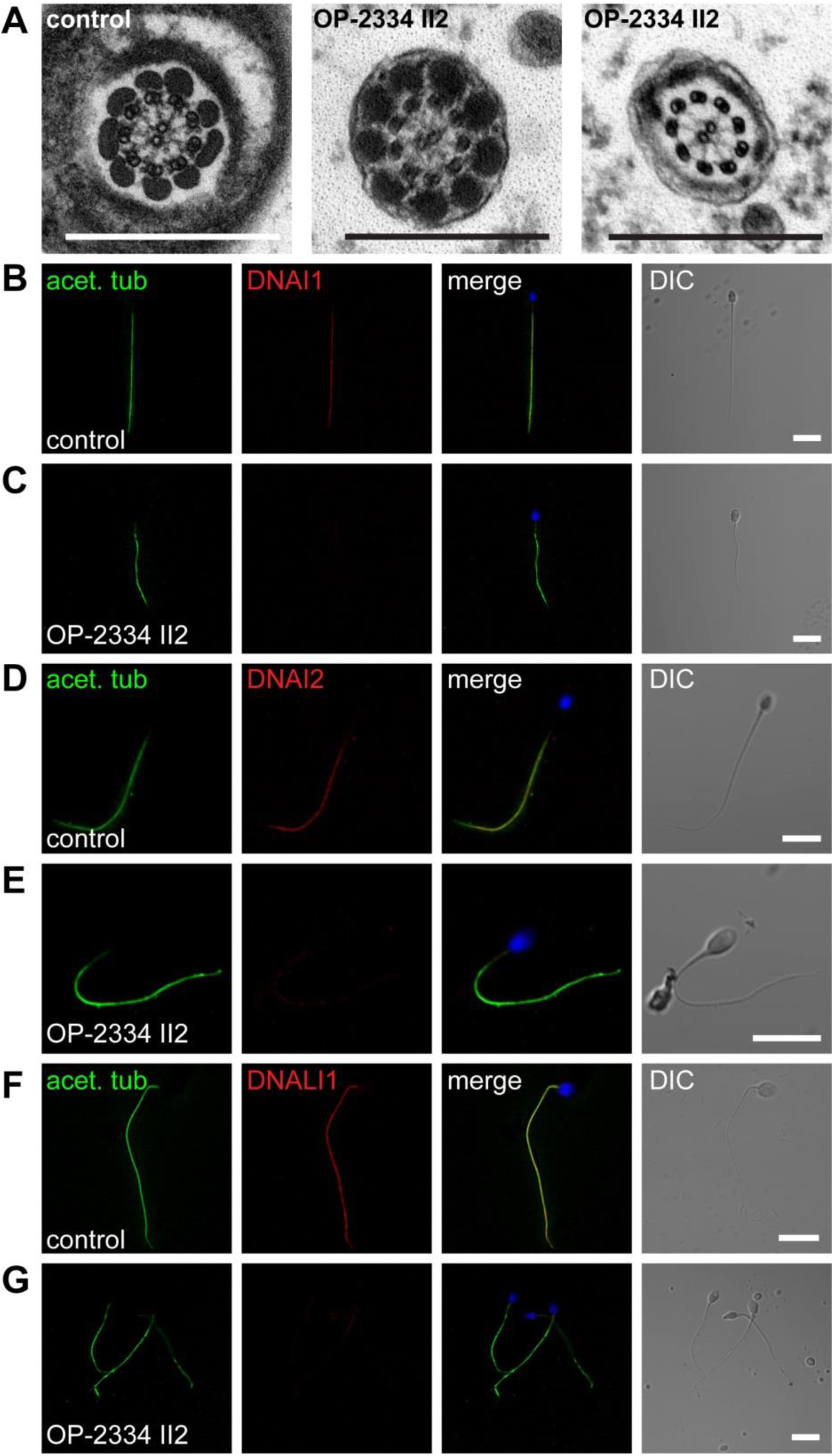
C11ORF70 is necessary for correct assembly of outer and inner dynein arms in sperm flagella. (A) TEM photographs of cross-sections of sperm flagella from a healthy control individual and from OP-2334 II2. The outer and inner dynein arms are present in control sperm flagella but are absent in the sperm flagella of OP-2334 II2. Scale bar represents 500 nm. (B to G) To confirm the ultrastructural defects of the dynein arms, sperms from an unaffected control (B,D and F) and OP-2334 II2 (C,E and G) were double-labeled with antibodies directed against acetylated tubulin (green), and ODA proteins (DNAI1, DNAI2), or IDA protein DNALI1 (red). DNAI1, DNAI2 and DNALI1 colocalize with acetylated tubulin along the sperm flagellum from the unaffected control (B, D and F, yellow). In contrast, DNAI1, DNAI2 and DNALI1 are absent or severely reduced in *C11ORF70*-mutant sperm flagella (C, E and G). Nuclei were stained with Hoechst33342 (blue). Scale bars represent 10 μm.

In order to understand C11ORF70 function within the cytoplasmic dynein arm assembly process and to investigate possible interactions with other proteins encoded by genes causing PCD, we next performed a yeast two-hybrid (Y2H) screen. Using C11ORF70 as bait we tested putative interactions with other known preassembly factors, ODA and IDA components, ODA-DC proteins, proteins assumed to be involved in dynein arm transport (WDR69, IFT46) and others (Table S2). Direct interaction between C11ORF70 and possible interactors was tested as previously described^(50)^.

All cDNA clones were confirmed by sequence analysis and matched RefSeq gene accession numbers. The screen revealed possible direct interaction with two assembly factors, namely DNAAF2 and PIH1D3 as well as with IFT46 (Figure S12). Because PIH1D3 and IFT46 pBD clones were found to be autoactivating (*data not shown*) we excluded these as interactions and considered only DNAAF2 to interact physically with C11ORF70.

Here, we demonstrate that deficiency of *C11ORF70* results in defects of ODAs and IDAs and ciliary immotiliy, and consequently altered mucociliary clearance and PCD. In one male PCD individual sperm immotility was observed, which is caused by disrupted assembly of dynein arms in sperm flagella, similar to findings observed in *DNAAF2/K*TU mutant individuals^(41)^.

In summary, we identify recessive loss-of-function mutations in the open reading frame *C11ORF70* in five PCD individuals from five distinct families. Interestingly, high resolution IF analyses showed that loss of C11ORF70 results in absence of the ODAs type 1 and type 2. However, the localization of the β-heavy chain DNAH11 was not affected. We have demonstrated that axonemal DNAH11 assembly is independent from other axonemal dynein heavy chains such as DNAH5 and DNAH9 and that the cytoplasmic preassembly of DNAH11 is not dependent on the function of DNAAFs such as DNAAF1 and DNAAF2^(6)^. Here, we demonstrate that loss of function of *C11ORF70* results not only in defects of the axonemal assembly of IDAs of group I2 (shown by the axonemal absence of DNALI1) but also from IDAs of group I3 (shown by the axonemal loss of DNAH6), indicating that C11ORF70 is involved in the assembly of those two IDA groups. Additionally, we demonstrated that deficiency of C11ORF70 results in absence of ODAs and IDAs in sperm flagella associated with male infertility. Binary interaction with DNAAF2 indicates a critical role for C11ORF70 within the dynein arm assembly machinery. We were unable to determine the cellular localization of C11ORF70 due to the lack of the availability of specific antibodies. However, the observed defects of ODAs and IDAs resembles findings observed in ciliary axonemes from individuals harboring mutations in *DNAAF* genes such as *DNAAF1*^(39,40)^ and *DNAAF3*^(42)^. Additionally, the direct interaction with DNAAF2 indicates that C11ORF70 is probably a new cytoplasmic preassembly factor and/or might be involved in transport of dynein components to the ciliary axonemes.

## Acknowledgements

We are grateful to the patients and their family members whose cooperation made this study possible, and thank all referring physicians. We thank A. Dorißen, D. Ernst, S. Helms, M. Herting, A. Robbers, L. Schwiddessen, F.J. Seesing, M. Tekaat, K. Wohlgemuth and C. Westermann for excellent technical work. The authors would like to thank the Genome Aggregation Database and the groups that provided exome variant data for comparison. A full list of contributing groups can be found at http://gnomad.broadinstitute.org/. This work was supported by the Deutsche Forschungsgemeinschaft OM 6/7, OM6/8, OM6/9, OM6/10, OM6/11 (H.O.), OL450/1 (H.Ol.), and HJ7/1-1 (R.H.), by the IZKF Muenster to H.O. (Om2/009/12 and Om/015/16), the European Union seventh FP under GA Nr. 305404, project BESTCILIA (H.O., K.G.N., M.C.P.) and “Innovative Medical Research” of the University of Muenster Medical School to N.T.L. (I-LO121517) and to J.W. (IWA121418). M.S. acknowledges funding from Radboudumc and RIMLS Nijmegen (Hypatia tenure track fellowship), the “Deutsche Forschungsgemeinschaft” (DFG CRC1140 KIDGEM) and the European research Council (ERC StG TREATCilia, grant No 716344).

## Web resources

The URLs for web resources used are as follows:

EMBL EBI Expression Atlas, http://www.ebi.ac.uk/gxa/home

Human Protein Atlas, http://www.proteinatlas.org/

Online Mendelian Inheritance in Man (OMIM), http://omim.org/

Varbank analysis software: https://varbank.ccg.uni-koeln.de/

Genome Aggregation Database (gnomAD), Cambridge, MA (URL: http://gnomad.broadinstitute.org)

## Description of supplemental data

Supplemental data includes Supplemental Material and Methods, 2 Tables, 12 Figures and 7 Videos

**Table S1.** Spermiograms from OP-2334 II2 show a severe ashenozoospermia (complete immotility) and a moderate oligospermia (reduced number of sperms).

**Table S2.** pAD-proteins tested by Y2H for direct interaction with pBD-C11orf70

**Figure S1.** Pedigrees of the PCD families with mutations in *C11ORF70* and sequence files.

**Figure S2.** Differentiation-dependent expression profiles of dynein arm preassembly factors in ALI-cultured nasal epithelial cells.

**Figure S3.** In situ hybridization analyses of wildtype mouse embryos reveal expression of 9230110C19Rik (*C11orf70*) exclusively at the left/right organizer.

**Figure S4.** *C11ORF70*-mutant respiratory cilia are deficient for the outer dynein heavy chain DNAH5. **Figure S5.** Absence of the outer dynein arm heavy chain DNAH9 in respiratory cilia of PCD individuals carrying *C11ORF70* mutations.

**Figure S6.** Loss-of-function *C11ORF70*-mutations result in absence of the outer dynein arm intermediate chain DNAI1.

**Figure S7.** Loss-of-function *C11ORF70*-mutations result in absence of the outer dynein arm intermediate chain DNAI2.

**Figure S8.** In *C11ORF70*-mutant respiratory cilia axonemal assembly of the outer dynein arm heavy chain DNAH11 is not affected.

**Figure S9.** In *C11ORF70*-mutant respiratory cilia the outer dynein arm docking complex (ODA-DC) is not affected.

**Figure S10.** Inner dynein arm light chain DNALI1 (IDA-Group I2) is absent in respiratory cilia of PCD individuals carrying *C11ORF70* mutations.

**Figure S11.** In *C11ORF70*-mutant respiratory cilia the inner dynein arm heavy chain DNAH6 (IDA-Group I3) is absent.

**Figure S12.** Identification of direct interaction partners of C11ORF70

**Supplemental Video 1.** High-speed videomicroscopy of *C11ORF70*-mutant respiratory cilia (OP-1416 II1)

**Supplemental Video 2.** High-speed videomicroscopy of *C11ORF70*-mutant respiratory cilia (OP-2249 II1)

**Supplemental Video 3.** High-speed videomicroscopy of *C11ORF70*-mutant respiratory cilia (OP-2334 II2)

**Supplemental Video 4.** High-speed videomicroscopy of control respiratory cilia

**Supplemental Video 5.** High-speed videomicroscopy of control sperm cells

**Supplemental Video 6.** High-speed videomicroscopy of *C11ORF70-*mutant sperm cells (OP-2334 II2)

**Supplemental Video 7.** High-speed videomicroscopy of *C11ORF70-*mutant sperm cells (OP-2334 II2)

